# Unravelling Genomic Origins of Lumpy Skin Disease Virus in Recent Outbreaks

**DOI:** 10.1101/2023.08.25.554819

**Authors:** Priya Yadav, Ankeet Kumar, Sujith S Nath, Geetha Shashidhar, Madhvi Joshi, Apurva Puvar, Sonal Sharma, Janvi Raval, Rameshchandra Pandit, Priyank Chavada, Sudheep Nagaraj, Yogesharadhya Revanaiah, Deepak Patil, S K Raval, Jigar Raval, Amit Kanani, Falguni Thakar, Naveen Kumar, Gundallhalli Bayyappa Manjunatha Reddy, Chaitanya Joshi, Baldev Raj Gulati, Utpal Tatu

**Author notes:** Corresponding author: Utpal Tatu.

## Abstract

Lumpy skin disease virus (LSDV) belongs to the genus *Capripox* virus and family *Poxviridae*. LSDV was endemic in most of Africa, Middle east and Turkey, but since 2015, several outbreaks have been reported in Asian countries. In this study we used Whole Genome Sequence (WGS) approach to investigate the origin of the outbreak and understand the genomic landscape of the virus. Our study showed that the LSDV strain of 2022 outbreak exhibited many genetic variations, compared to the Reference Neethling strain sequence and the previous field strains from India. A total of 1819 variations were found in 22 genome sequences, which includes 399 extragenic mutations, 153 insertion frameshift mutations, 234 deletion frameshift mutations, 271 Single nucleotide polymorphisms (SNPs) and 762 silent SNPs. 38 genes have more than 2 variations per gene and these genes belong to viral-core protein, viral binding proteins, replication and RNA polymerase proteins. We highlight the importance of several SNPs in various genes which may play an essential role in pathogenesis of LSDV. Phylogenetic analysis performed on all whole genome sequences of LSDV showed two types of variants in India. One group of the variant with fewer mutations was found to lie closer to the LSDV 2019 strain from Ranchi while the other group clustered with previous Russian outbreaks from 2015. Our study highlights the importance of genomic characterization of viral outbreaks to not only monitor the frequency of mutations but also address its role in pathogenesis of LSDV as the outbreak continues.

## INTRODUCTION

The Lumpy Skin Disease is caused by Lumpy Skin Disease Virus (LSDV) [1] is a double-stranded DNA virus of genome size 150 kb and belongs to the genus *Capripoxvirus*, Sub-family *Chordopoxviridae* and family *Poxviridae*[2]. The other members of the genus are known as Goat poxvirus (GTPV) and Sheep poxvirus (SPPV). LSDV is very similar to all the members of the Poxvirdae family in morphological characteristics such as it is very similar to the vaccinia virus under Electron microscopy [3]. LSDV is non-zoonotic and known to infect specific hosts including cattle (*Bos indicus*, *Bos taurus*) and domestic water buffaloes (*Bubalus bubalis*) [4–6]. In addition, LSDV can also infect mammals such as camel, giraffe, wildebeest in the wild [7–9]. It spreads via direct contact through skin lesion, milk as well as blood-sucking insects such as biting flies, ticks and mosquitoes [10–13]. The infection spreads more in warm and wet period as compared to the winters due to increased insect population and its mobility in summers [14]. The virus generally spreads from an infected animal to a healthy animal via viral shedding through skin lesions, nasal discharge, saliva, and lachrymal secretion [11,15].

The Lumpy skin disease virus was first reported in 2019 from India and has since caused severe outbreaks. In the recent outbreak, cases started to appear since May 2022. Almost all the states reported cases, but 15 Indian states experienced a significant economic loss with a death toll nearing 1,00,000 cattle [16]. Since in a developing country like India, livestock production constitutes one of the important ways of earning a livelihood, a deadly disease such as Lumpy Skin Disease has caused direct loss to the economy and poor production of livestock [16]. India has a cattle population of 308 million; therefore, controlling the spread of infectious diseases is important [17].The direct loss includes dead cattle, a decrease in milk production while the indirect losses include movement restriction of cattle across the country [16]. Earlier studies have reported pathological changes to most of the organs and tissue of infected animals such as cow mastitis, necrotic hepatitis, lymphadenitis, orchitis and also in some cases, myocardial damage [18].

WHO has now identified lumpy skin disease as a notifiable disease [19]. Since its first discovery in Zambia in 1931 [20], this disease was initially confined to the Sub-African region until 1989 and then it started spreading across boundary to Middle East Asia [21,22]. LSDV was reported in 2016 in Russia and other South-East European nations [22]. The disease first appeared in India along with other Asian countries such as China, Nepal, Thailand, Bangladesh, Bhutan in November 2019 [23,24]. While LSD has been reported in India since 2019, it has caused significant damage during 2022, infecting more than 2 million cattle. The symptoms of LSD vary in individual animals depending on the severity of the infection. An animal takes 1-4 weeks to develop symptoms such as high fever, ocular and nasal discharge, loss of appetite, nodular lesion on skin [25]. According to the data available, a mortality rate of 5-45% was observed [26–29]. The major states affected in terms of mortality and morbidity are Rajasthan, Gujarat, Uttar Pradesh, Punjab, Haryana, Karnataka, West Bengal and Maharashtra [30]. The Indian government has taken several measures to control the spread of LSD, including mass vaccination campaigns, setting up quarantine facilities and restricting the movement of infected and susceptible animals. However, the disease continues to be a challenge due to a lack of awareness about its transmission and control, as well as the difficulty in detecting infected animals in the early stages of the disease.

It is generally recognized that the LSDV may have originated from one of the earlier pox virus species and then evolved by spreading in different kind of hosts. Double-stranded DNA viruses are known to use homologous recombination for evolution towards expanding their host range and virulence [31]. In this study, we have utilized genome sequencing to identify the variants of LSDV circulating in India. Through the phylogenetic analysis we found there are two different classes of variants in India. We then performed mutation (SNP) analysis and found the groups differ significantly in the number of mutations.

## MATERIAL AND METHOD

### Sample collection

Biological specimens including saliva swab, blood sample, skin lesion scabs were collected from infected cattle as per the standard practices without using any anaesthesia by veterinarians in Karnataka in India. A due consent was taken from the cattle owner before collection of samples. The animals presented symptoms such as fever, nasal discharge and characteristic pox nodular skin lesions. Blood samples were collected in EDTA vaccutainers (Becton-Dickenson) and the skin lesions were collected and transported to the laboratory in 50% glycerol. The samples were used to extract DNA and remaining samples were stored in −80°C for further use.

### Processing of Samples

For molecular diagnosis, 200µl of Blood sample was used to extract DNA by using Huwel Nucleic acid extraction kit (HL-NA-100) and the DNA concentration was estimated using nanodrop spectrophotometer.

For the present study, total 22 samples of LSDV were used for sequencing. 7 samples were from the outbreaks during 2020-2021, 1 sample from Ranchi outbreak in 2019 and 14 samples were from 2022 outbreak in different parts of India, as mentioned in the table below. Out of total 22 samples, 10 viral samples were taken from cell-infected viral cultures while 12 samples were taken from skin nodules from cattle.

Sample metadata:

| Sl. No. | State | District | Year | Species | Age | Source of virus | Sample ID |
| --- | --- | --- | --- | --- | --- | --- | --- |
| 1. | Ranchi | Ranchi | 2019 | Cattle | Adult | Cell infected culture | Ranchi 2019 L1 |
| 2. | Rajasthan | Banswara | 2021 | Cattle | Adult | Cell infected culture | Banswara 2021 L2 |
| 3. | Gujarat | Surat | 2021 | Cattle | Adult | Scab | 1 Surat 2021 N6 |
| 4. | Gujarat | Surat | 2021 | Cattle | Adult | Scab | 2 Surat 2021 N7 |
| 5. | Gujarat | Jamnagar | 2022 | Cattle | Adult | Scab | 1 Jamnagar 2022 N1 |
| 6. | Gujarat | Jamnagar | 2022 | Cattle | Adult | Scab | 2 Jamnagar 2022 N2 |
| 7. | Gujarat | Jamnagar | 2022 | Cattle | Adult | Scab | 3 Jamnagar 2022 N5 |
| 8. | Gujarat | Anand | 2022 | Cattle | Adult | Scab | 1 Anand 2022 N8 |
| 9. | Gujarat | Anand | 2022 | Cattle | Adult | Scab | 2 Anand 2022 N9 |
| 10. | Rajasthan | Bikaner | 2022 | Camel | Adult | Cell infected culture | Camel 2022 L3 |
| 11. | Rajasthan | Jalore | 2022 | Cattle | Adult | Cell infected culture | Jalore 2022 L4 |
| 12. | Rajasthan | Nohar | 2022 | Cattle | Adult | Cell infected culture | Nohar 2022 L5 |
| 13. | Maharashtra |  | 2022 | Cattle | Adult | Scab | MH01 2022 |
| 14. | Maharashtra |  | 2022 | Cattle | Adult | Scab | MH02 2022 |
| 15. | Maharashtra |  | 2022 | Cattle | Adult | Scab | MH03 2022 |
| 16. | Maharashtra |  | 2022 | Cattle | Adult | Scab | MH04 2022 |
| 17. | Karnataka | Raichur | 2020 | Cattle | Adult | Cell infected culture | KA05 2020 |
| 18. | Karnataka | Bengaluru | 2020 | Cattle | Adult | Cell infected culture | KA06 2020 |
| 19. | Karnataka | Ramanagara | 2021 | Cattle | Adult | Cell infected culture | KA07 2021 |
| 20. | Karnataka | Haveri | 2021 | Cattle | Adult | Cell infected culture | KA08 2021 |
| 21. | Karnataka | Maddur | 2022 | Cattle | Adult | Cell infected culture | KA09 2022 |
| 22. | West Bengal |  | 2022 | Cattle | Adult | Scab | WB 2022 |

### Molecular Diagnostics

The primers were designed for the viral Fusion gene to confirm LSDV in the collected samples, forward primer-5’ ATGGACAGAGCTTTATCA, reverse primer-5’ TCATAGTGTTGTACTTCG (10pM/µl) with Tm 55°C. The conventional PCR was carried out in 25µl reaction with the following conditions used: 94°C for 5 mins for initial denaturation, 30 cycles for 94°C for 1 min denaturation, 55°C for 30 sec annealing, 72°C for 45 sec extention and a final extention step of 72°C for 5 mins. The PCR result was visualized on 1% agarose gel (Himedia) having Ethidium Bromide and analysed against 1Kbp DNA ladder (Takara) in (1X) TAE buffer[32].

### Individual PCR for Sequencing

A total of 55 overlapping primer sets for 3 kbp amplicons with Tm 66°C were designed by using PrimalScheme [33], a tool which is used to design primers for multiplex PCR, especially for viral outbreak strains. The complete genome sequence of LSDV from previous outbreak with GenBank ID: OK422493.1 was used to design primers. The DNA extracted from the skin scab sample collected from India was used to amplify the LSDV DNA. The 2-step PCR was performed using Phusion polymerase with the following conditions: 98°C for initial denaturation, 98°C for denaturation, 66°C for annealing and extension, and 72°C for 3 mins for the final extension step. The PCR results were analysed on 1% agarose gel (Himedia) having Ethidium Bromide and analysed against 100 bp DNA ladder.

### Sequencing using Oxford-Nanopore Technology

The amplified LSDV DNA amplicons of 3 kbp size were pooled together, and the DNA concentration was determined by using Qubit 3 fluorometer (Invitrogen) after calibrating with the standards. The samples were processed for genome sequencing on MinION (Oxford Nanopore Technologies, Oxford, United Kingdom) with the ligation sequencing kit 1D, SQK-LSK109 (Oxford Nanopore Technologies). Samples were sequenced on flow cell R10.3 Version (FLO-MIN111) with 1570 active pores. For the MinION, the Barcode 01-09 from the native barcoding kit (Oxford Nanopore Technologies) were used. The DNA was cleaned up by using KAPA Hyperpure beads (Roche) after every step during sample preparation and DNA concentration was recorded. The DNA library was prepared as per the Oxford Nanotechnologies protocol by adapting the following steps: end prep of the LSDV amplicons followed by Barcoding and adapter ligation [34].

### Sequencing using Illumina NGS Technology

NGS library preparation for LSDV whole genome sequencing was done using in solution Tagmentation based Illumina Nextera XT DNA Library Prep kit with starting DNA concentration of 1 ng/µl. Prepared libraries were quantified using Qubit 1X ds DNA HS kit and quality check was done on Agilent 2100 bioanalyzer using High sensitivity DNA reagent kit. All the libraries were normalized up to 4 nM and pooled. The equimolarized pool of library was processed for the denaturation with 1% PhiX library spike-in as control library. Sequencing was performed on Illumina NovaSeq 6000 SP Reagent Kit v1.5 (300 cycles) paired-end chemistry. 5 Isolated Viral DNA library was sequenced on Illumina MiSeq using MiSeq reagent kit V2 (500 Cycles).

### MinION sequencing data analysis

The flow cell was primed by using EXP-FLP002 Flow cell priming kit. 300 fmol of DNA library was loaded along with the sequencing buffer and loading beads on the SpotON MinION sequencer and was run for 11 hours using a new R10.3 flow cell. Basecalling and demultiplexing reads by barcode was performed by using Guppy command-line software. Primer scheme was used in Artic pipeline and primers were trimmed and reads were aligned to the LSDV whole genome sequence GenBank ID: OK422493 from Ranchi, India in 2019. Alignment-based consensus were generated using protocol adapted using Artic pipeline developed for SARS-COV-2 and the gaps were filled by amplifying the LSDV again and the same protocol was followed for sequencing. The generated sequence was annotated into genes and proteins by using GeneMarkS2 [35], which covered 156 genes in the generated LSDV sequence.

### Phylogenetic analysis and SNP analysis

The newly generated consensus sequence was used along with other 65 LSDV complete genomes and Goat poxvirus sequence retrieved from the public domain database Bacterial and Viral Bioinformatics Center (BV-BRC). A Bayesian phylogenetic tree was constructed using 1000 Bootstrap iterations using MAFFT. The aligned sequences were used to build phylogenetic tree using MEGA XI. The Reference Neethling strain genome sequence was used to align the generated LSDV 2022 India genome sequence and nucleotide mutations were analysed. R script was used to detect SNPs.

### Ethics Statement

Institute Bio-Safety certificate (IBSC) with Reference number IBSC/IISc/UT/16/2023 was obtained from the ethics committee to work on LSDV as it is non-zoonotic. The LSDV samples were collected by the veterinarians as per the approved protocols and guidelines.

## RESULTS

### Analysis of LSDV cases in the field reveal varied signs and clinical outcomes

Samples of blood, swab and samples of skin scab from infected cattle were collected in Karnataka, Maharashtra, Rajasthan, Jhansi during August-November 2022 while the outbreak was going on in India. The animals were showing typical clinical signs such as reduced milk yield, loss of body mass, raised body temperature upto 40-41°C, nasal and lachrymal discharge, lost appetite, and skin nodules observed on the body of both young and adult cattle. The skin nodules burst open when the viral titre is high, exposing the internal layers of the skin which leads to open wounds and eventually secondary bacterial infection if not timely treated (Figure-1A). The villages observed mortality rate as high as 40% for infected cattle, though higher mortality was evidenced in young cattle as compared to the adults.

Vectors such as flies, blood-sucking ticks, and mosquitoes were also observed in the vicinity of the animals. Infected animals shed the virus through nasal discharge, lachrymal fluid, and saliva, facilitating the easy spread of LSDV in close proximity settings. Due to the lack of proper ventilation and isolation facilities for cattle, LSDV managed to infect over 20 million cattle in 15 different states, and the outbreak is still ongoing. The 2022 outbreak in India has caused significant economic losses, as the country heavily relies on agriculture and farming, which employs 50% of the population.

### PCR-based detection strategy for LSDV

The diagnosis for Lumpy skin disease mainly depends on the typical clinical signs and differential diagnosis. The early signs of LSDV overlaps with other diseases like foot and mouth disease or insect bites. Agar gel precipitation is also used to detect viral antigen in a serum or tissue sample. But this test is not specific and cannot be used for LSDV because the antigens of LSDV are shared with other poxviruses. Therefore, molecular diagnosis by using PCR is the most effective method to detect LSDV.

Samples including blood, oral swab, skin scabs were collected from different parts of India and transported to the ICAR and IISc laboratory to confirm LSDV infection. Conserved regions of the virus genome sequence were identified across different strains and used as a target for detection by multiple sequence alignment. LSDV specific primers were used for fusion gene LSDV0117, which helps the viral envelope fusion with the host membrane as described in method section. Conventional PCR was performed and the amplicon size of 472 bp was confirmed (Figure 1B). All the samples collected from symptomatic cattle were positive for Lumpy Skin Disease Virus. The DNA extracted from blood and skin scab gave good amplification as compared to the DNA extracted from the mouth swab.

**Figure 1.**
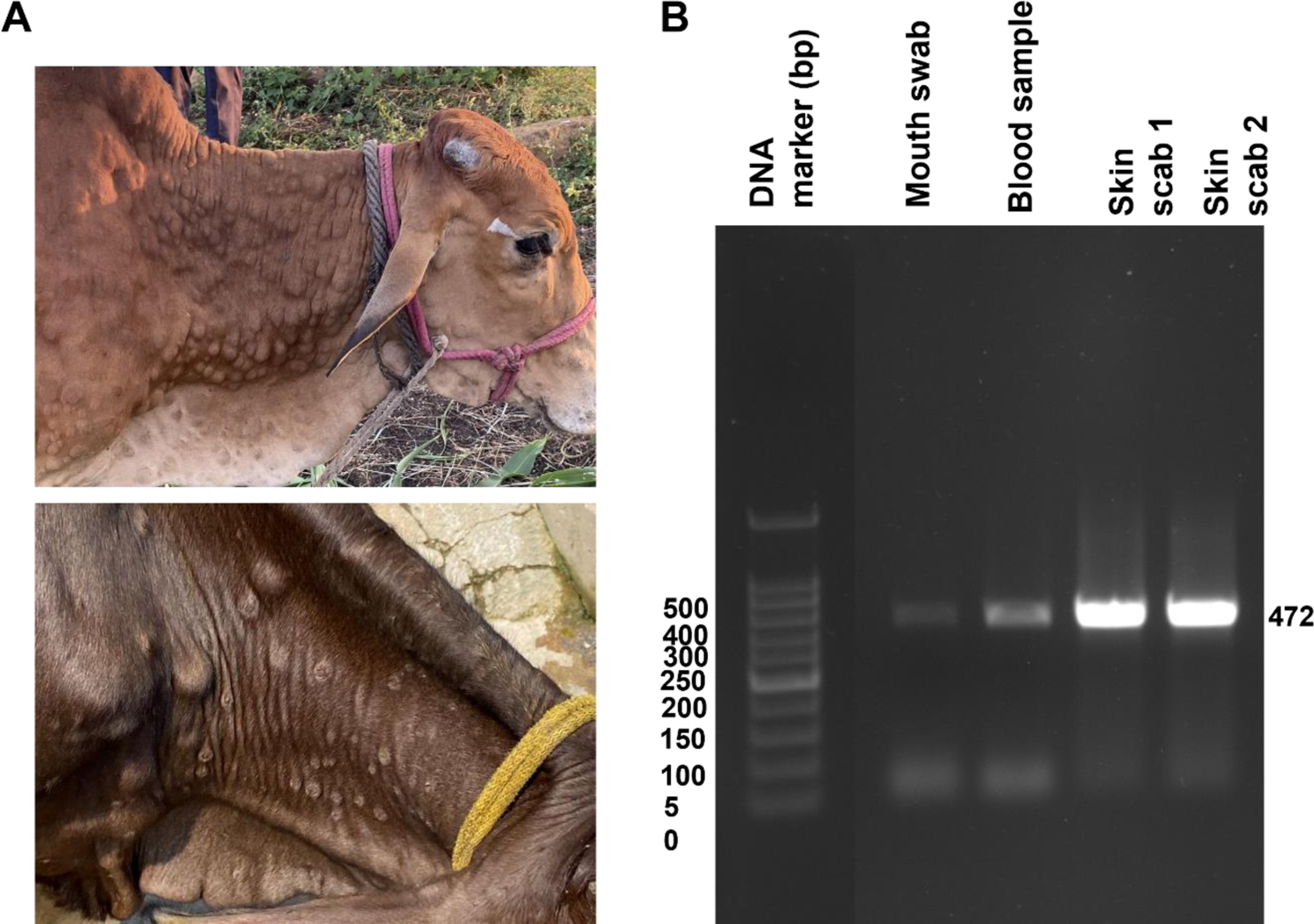
Clinical Signs and molecular diagnostics. A) Animal showing severe skin lesions and nodules on body and appearance of swollen lymph nodes B) PCR-based confirmation of LSDV by using partial viral fusion gene amplification.

### Amplification of Indian LSDV strain genome to develop amplicon-based whole genome sequencing

DNA extracted from 22 samples (including twelve skin scab/nodule samples from cattle and ten virus samples collected after cell culture passage) was used for further processing. The whole genome of LSDV is approximately 150 kb, so it was necessary to generate smaller amplicons (3 kb) to achieve full genome coverage for Oxford Nanopore technology and overlapping primers were designed. Nanopore sequencing produces long reads, which are useful for characterizing complex genomic regions.

A total of 1,126,683 reads were obtained, ranging in size from 105 kb to 54,253 kb, with a median size of 10.686 kb. Reads smaller than 1.5 kb were filtered out to avoid non-specific reads. The coding sequences (CDS) of 156 genes were obtained using GeneMarkS2 for *ab initio* gene prediction. The genome was assembled from the raw sequencing data, and the location of the genes was identified. Additional 12 samples collected from Jamnagar, Anand, and Surat were sequenced using Illumina sequencing, covering over 90% of the genome sequence. To date, there are 56 whole genome sequences of LSDV available in public databases. Therefore, our study, which includes the sequencing of an additional 22 LSDV genomes, is highly relevant for monitoring circulating strains and developing vaccine strategies.

### Phylogenetic analysis of whole genome sequences of LSDV reveals multiple strains

Multiple sequence alignment of the LSDV whole genome sequences from different regions of the world was performed and this information was used to construct a maximum likelihood-based phylogenetic tree, which shows the evolutionary relationships among the different LSDV strains. Goat poxvirus, Goat poxvirus vaccine (GV1) and Sheep poxvirus vaccine (SV1) were used for comparison, as shown in Figure 2.

**Figure 2.**
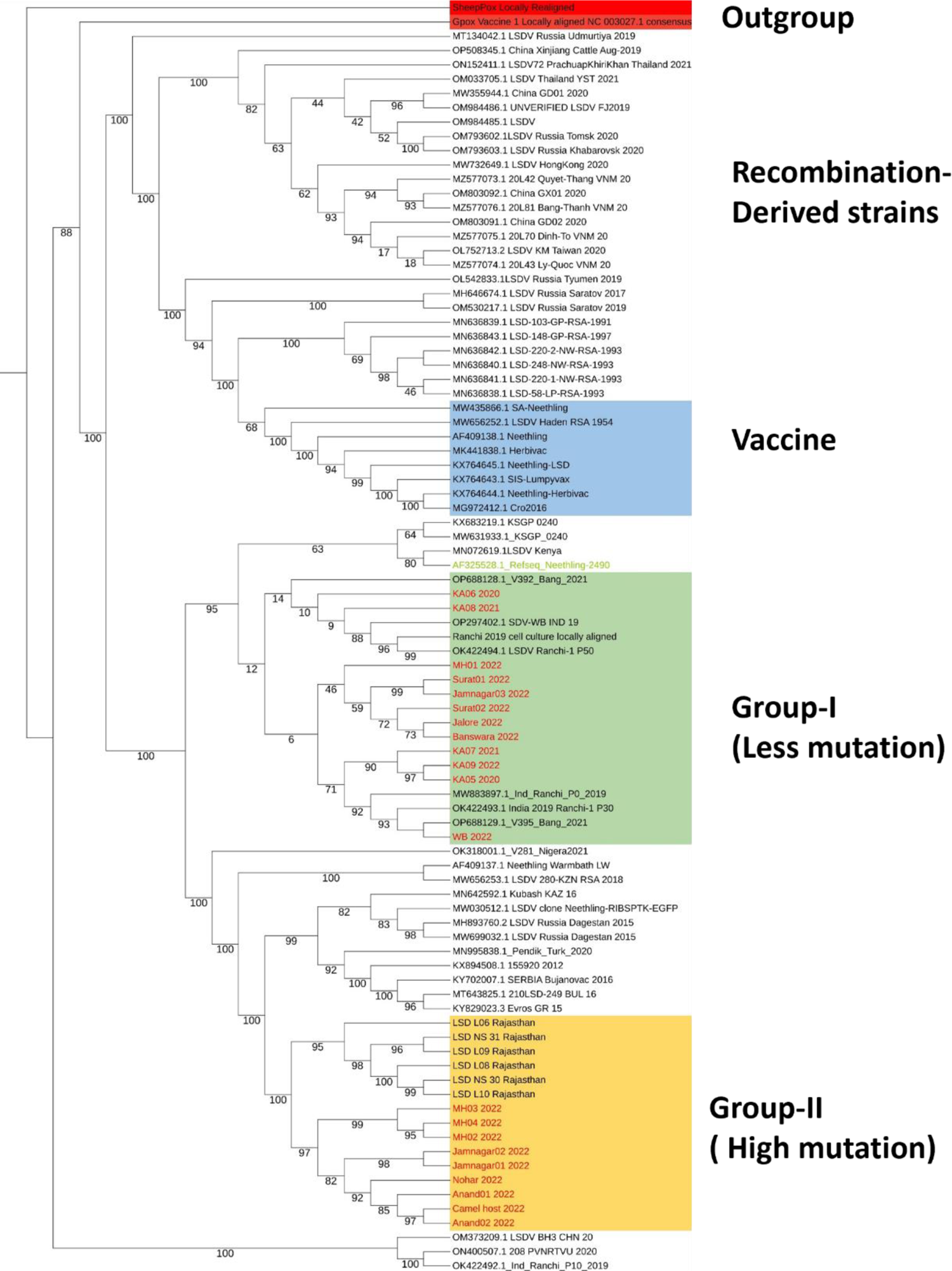
Phylogenetic analysis of LSDV. Bayesian phylogenetic tree based on whole genome sequences showing the relationship in Capripox virus family including Sheep poxvirus, Goat poxvirus and complete genome sequences of LSDV strains sequenced from many parts of India during 2022 outbreak as well as sequences available on NCBI from previous outbreaks. The genomes sequenced in this study are highlighted in red text.

The sequenced genomes from 2022 outbreak in India lies on two separate branches on the phylogenetic tree, which has been named as Group I and Group II (Figure 2). Group I genomes closely resemble the Ranchi sequence from 2019 outbreak and Hyderabad sequence from 2020 outbreak. This group has lesser number of variations in their genome sequences as compared to the reference Neethling LSDV sequence. The other group, Group II, with higher number of variations per sequence, lies separately, grouping with Russian 2015 LSDV outbreaks. This observation suggests that the current 2022 LSDV outbreak in India is a result of two different group of strains circulating together in the same region.

There are five main branches on the phylogenetic tree. The Indian LSDV sequences from 2022 are distinct from the vaccine strains of LSDV, confirming that the recent outbreak in India is unlikely to be the result of vaccine spillover. All the Neethling virus-based vaccine strains and vaccine-derived recombinant strains, characterized by combined genetic sequences [36], form a separate group. The close resemblance of recent outbreak strains in Group I to the 2019 Ranchi strain suggests that infections were occurring, although not very severe, in cattle for a few years. However, the higher number of mutations in Group II indicates the circulation of another strain with an increased mutation rate in genes involved in host cell binding and immune evasion.

The Indian LSDV sequences are 99.98% identical to the reference Neethling LSDV virus and 97.38 % similarity with Goat poxvirus. Upon multiple sequence alignment of Goat poxvirus with LSDV sequences, it was found that around 3% of the whole genome is not identical, making it an outgroup and most of the differences lies in the end terminal repeat region of Goat poxvirus. This end terminal region consists mainly of genes important for virulent factors such as ankyrin repeat-domain containing proteins which helps the virus to bind to the host cell. The resulting phylogenetic analysis is important for understanding the evolution of the virus and for developing effective control strategies to limit its spread. It also provides valuable information for monitoring and detecting new outbreaks and for guiding the development of appropriate vaccines and therapeutic interventions.

### SNP analysis in the genome sequence reveals genetic variants with potential to cause severe infection

We performed mutation analysis using nucmer and found a total of 1819 variations in the 22 sequenced genomes compared to the reference sequence LSDV Neethling strain (AF325528). These variations at the amino acid level were dominated by silent SNPs, followed by extragenic mutations, SNPs, deletion frameshift mutations, and insertion frameshift mutations (Figure 3A). Group II sequences exhibit the highest number of SNPs at the amino acid level (Figure 3B). Among the LSDV genes, 38 genes have more than two variations per gene, with the gene encoding Interleukin-10-like protein having the highest number of variations (10). This is followed by the virion core protein gene with 7 variations, DNA ligase-like protein, B22R-like protein, and Ankyrin-like protein with 6 and 5 variations, respectively (Figure 4). Other genes, such as Kelch-like proteins, RNA polymerase subunit, EV glycoprotein, DNA helicase transcriptional elongation factor, and early transcription factor large subunit, important for virus binding to host cells and virus replication, have three to four variations in each gene. Additionally, 25 genes have two mutations, including several hypothetical protein coding genes, and 34 genes show one variation only.

**Figure 3.**
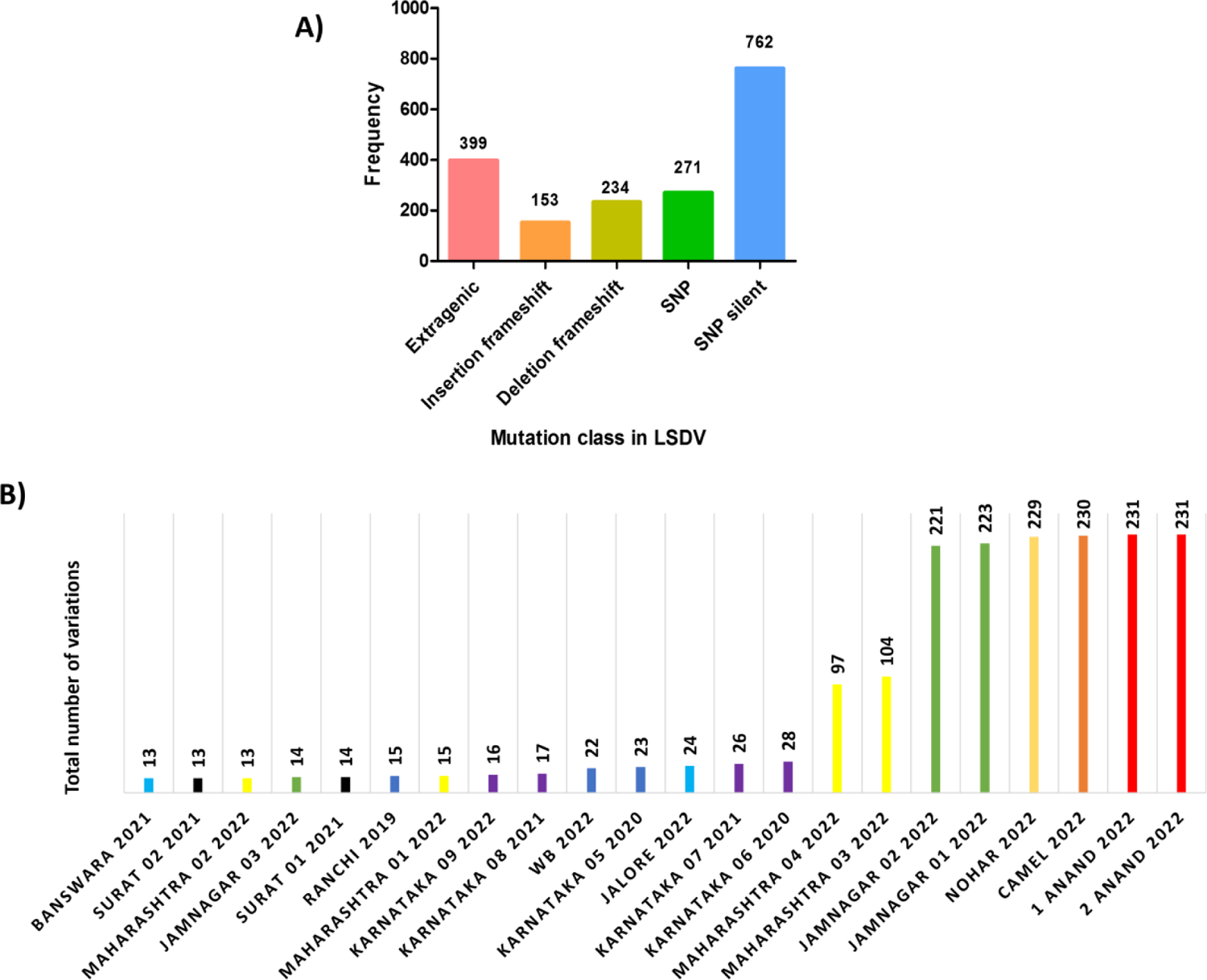
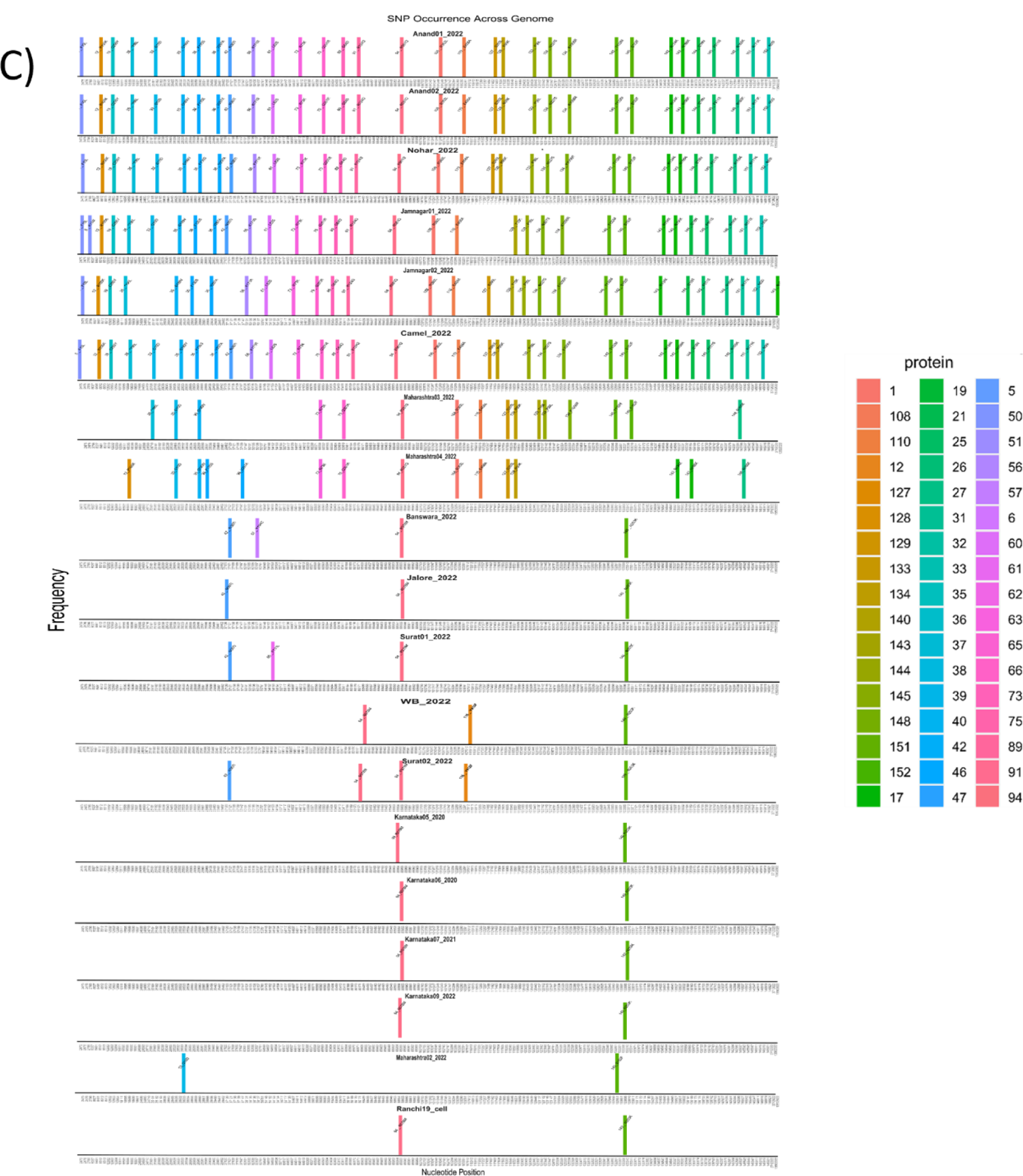
The variations in 22 LSDV genome sequences from India were classified into mutation classes. A) All classes of variations present in a total of 22 genomes, most of the variations belong to SNP_Silent followed by extragenic mutations B) Total number of variations identified in LSDV strains from 2022 outbreak. The colors in the graph used are for various regions in India C) Single Nucleotide Polymorphisms (SNPs) on amino acid level across all LSDV genomes sequenced.

**Figure 4.**
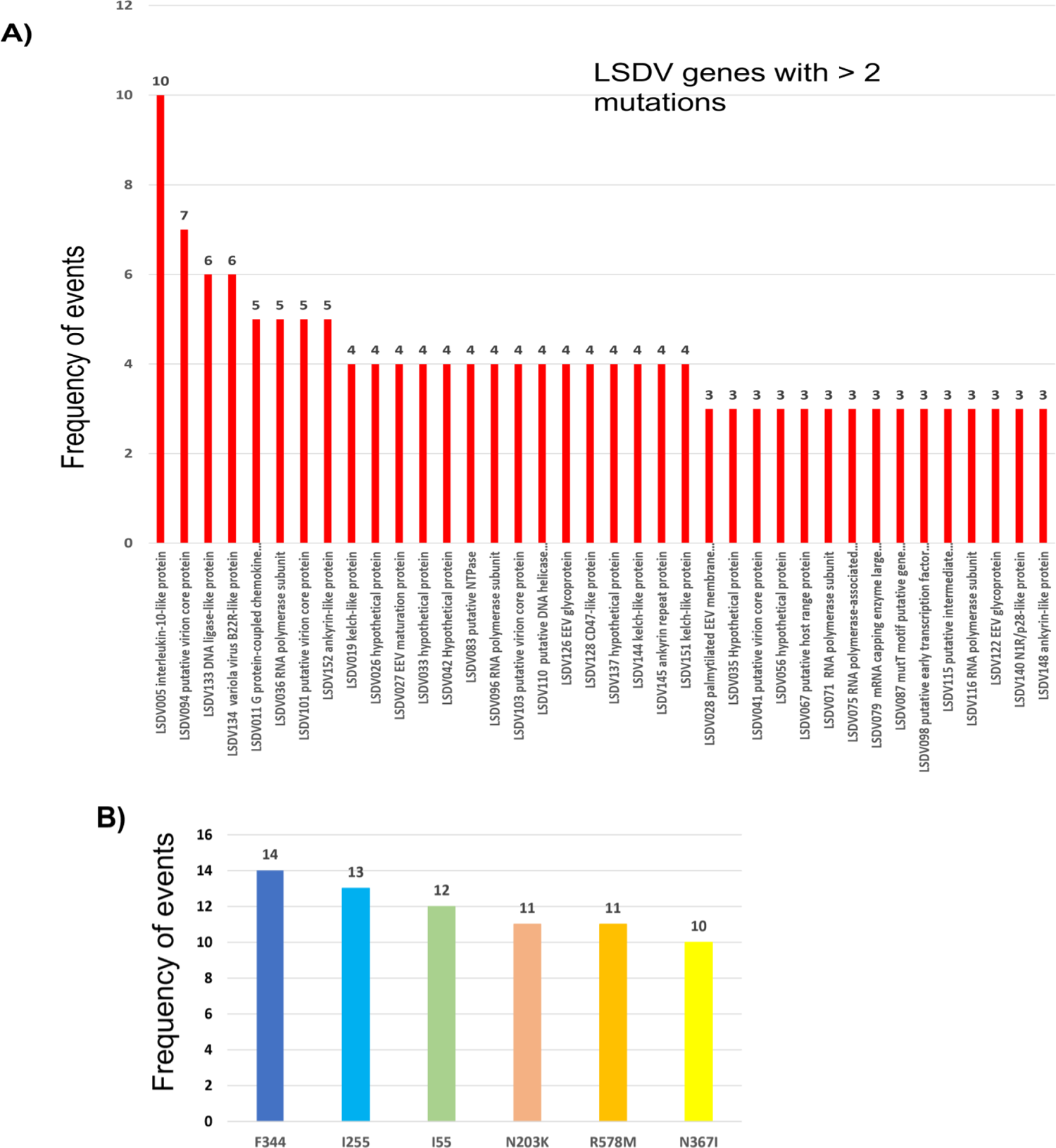
A) Graph showing number of variations in protein-coding genes in LSDV, with more than 2 variations per gene represented. B) The graph represents the most common mutation on amino acid level in 22 LSDV genome sequences upon analysis.

The most mutated protein coding gene, namely LSDV005, coding for Interleukin-10-like protein is a functional viral cytokine homolog that plays a role in regulating host immune response, as inferred from Uniprot. This protein has 4 helix cytokine-like core. Viral interleukins have been shown to activate cellular signalling cascades that enhance viral replication [37]. Previous gene knockout studies have shown that LSDV005 is one of the important protein coding gene which is responsible for virulence of LSD virus in host cells. LSDV005 is similar to cellular IL-10 in the carboxyl terminus part and affects the immune system similarly [38].

Another highly mutated gene is LSDV094 which codes for virion core protein. LSDV genome consists of 10 virion core protein coding genes throughout the genome which make up the core/nucleoprotein of the LSD virus. One common SNP in this gene in Group IIb leads to R578M transition, that is from basic to non-polar amino acid. Other variations in this gene are F111F, S168S, F354F, R581Q, I635I. Virion core proteins accounts for the one of the largest family of poxvirus proteins, many of them have been correlated as virulence factors [39].

Other genes with major variations include LSDV134 which encodes for variola virus (VV) B22R like protein. It is a VV immunomodulating gene with sequence homology to serine protease inhibitors (serpins) that possess antiapoptotic and anti-inflammatory properties. B22R has been shown to reduce host’s immune response to the virus and rabbitpox equivalent of VV B22R has been shown to inhibit apoptosis in a caspase-independent manner and increase host range as well [40].

Another important gene which is mutated on protein level is LSDV036 which encode for RNA polymerase subunit. Poxviruses encode a multi-subunit DNA-dependent RNA-polymerase that carries out viral gene expression in host cytoplasm. There were two silent mutations, and 2 SNPs observed in Group II genome sequences. The SNP leads to F130S, V297A substitution and given its role in the regulating transcription, any mutations in this gene can leads to altered ability to proliferate and cause symptoms.

Gene LSDV140 encoding for Ring finger host range protein or p28-like protein was also found to be mutated with N203K transition in Group I while Group II showed SNPs T132M and S152F. P28-like protein is a part of Ub system and act as Ubiquitin ligase. Similar mutation was reported earlier also in this protein [41]. These findings highlight the importance of consistent monitoring of genetic variations among LSDV variants. Rest of the SNPs belong to hypothetical proteins, whose functions are yet to be identified in poxviruses. Out of 156 genes of LSDV, more than 40 protein coding genes are hypothetical, and their function remains unknown. 26 of these hypothetical protein coding genes were found to have variations. There is a total of 399 extragenic mutations present in 22 genome sequences and most of the variations were observed in 3’UTR. Since 3’UTRs are involved in post-transcriptional regulation, and mRNA stability and degradation, these variations might affect gene expression and regulation in the host cell.

Many unique mutations were also identified in the genomes sequenced in the current study, such as N1 sample collected from Jamnagar, Gujarat in 2022 has K114I in gene LSDV005 coding for Interleukin-10-like protein; West Bengal sample showed unique SNP Y418F in gene LSDV116 coding for RNA polymerase subunit; Banswara, Rajasthan 2021 genomes sequence showed Y194C in LSDV057 coding for putative virion core protein. Mutations in these proteins can be one of the sophisticated mechanisms for enhancing the spread of virus and these mutations can affect the virulence, host range and antigenicity of the virus. These mutations can arise spontaneously or due to selection pressure from the host immune system and from the use of vaccines.

The most common mutations found in Indian LSDV 2022 sequence include changes in the viral envelope protein, which can alter the ability of virus to evade the host immune response, or in the viral polymerase gene, which can affect the replication and spread of the virus. It is important to monitor the genetic variation in LSDV in India, as it can help to understand the evolution and transmission of the virus and to design effective control strategies, including vaccines and diagnostic tests.

## DISCUSSION

The frequency in the occurrence of the outbreak and spread of the virus from its original geographic range has led to increased research. The first major outbreak in India happened in 2019, most of the sequences deposited for that outbreak are closely related to Neethling reference strain showing a few mutations. But the recent outbreak in India shows presence of at least two types of variants circulating in India.

To gain a comprehensive understanding of the variants of LSDV and their circulation patterns, samples were collected from multiple parts of India. The phylogenetic analysis of these sequences revealed two major groups. Analysis of the sequence composition at the nucleotide and protein levels demonstrated significant differences in the occurrence of single nucleotide polymorphisms (SNPs) between the two groups. The group that is closer to the Neethling strain exhibited less variation, while the other group displayed a higher number of mutations, most of which did not result in amino acid changes. Generally, the pox viruses are not fast evolving viruses, such as Variola virus is estimated to be evolving at ∼ 0.9-1.2 × 10^-6^ substitutions/site/year [42]. But they can undergo mutations due to mechanisms such as homologous and nonhomologous recombination, gene duplications, gene loss, and the acquisition of new genes through horizontal gene transfer.

Recently, Schalkwyk et al. established the substitution rate of 7.4 × 10^-6^ substitutions/site/year in Lumpy skin disease virus [43]. Therefore, it is possible that the high severity in the recent outbreaks has increased due to the increased number of mutations, as seen in Group II, containing sequences from Indian regions with high severity cases such as Rajasthan, Gujarat, and Maharashtra, which enables the virus to replicate more and cause more clinical symptoms and resulting in high mortality. Similarity in mutations of Group II strains with Nigeria 2021 and Russian 2015 outbreak suggests that there might have been a transboundary migration of an infected animal as both groups show higher number of similar mutations. Some SNPs caused changes in amino acids within proteins, particularly in genes like Interleukin 10-like protein, virion core-like protein, and GPCR-like protein. These mutations could be due to the evolutionary pressure on the virus resulting from the widespread use of vaccines that are not very effective, leading to mutations in important viral immune response genes.

Field analysis of LSD cases revealed a wide range of clinical signs and outcomes in infected cattle, including reduced milk production, weight loss, elevated body temperature, nasal and lachrymal discharge, loss of appetite, and the development of skin nodules. When these nodules burst open, they can lead to open wounds and bacterial infections. The high mortality rate, especially among young cattle, highlights the severity of LSDV infection. The increased severity observed in samples from Gujarat, Maharashtra, and Rajasthan may be a result of the high number of variations in those regions. Analysis of LSDV genomes identified numerous genetic variations, including silent SNPs, extragenic mutations, deletion and insertion frameshift mutations, and SNPs leading to amino acid changes. Several genes with multiple variations were found, including those encoding Interleukin-10-like protein, virion core protein, RNA polymerase subunit, and B22R-like protein. These variations can influence virulence, host range, and virus antigenicity. Recently, there have been reports of LSDV infection in camels, freely-ranging gazelles in India, and giraffes in Vietnam [8,9,44], indicating that LSDV can undergo transmission across species with an increased number of mutations. A total of 230 variations were observed in the LSDV strain sequenced from camel hosts. These SNPs were similar to the strains in Group II sequenced from cattle hosts. Mutations in proteins like B22R, coded by LSDV134, which have been shown to reduce the host’s immune response to the virus and increase host range, were also found in the LSDV strain from camels [40]. Therefore, it becomes important to identify and vaccinate other hosts which can act as reservoirs for the virus. Monitoring genetic variations in LSDV is crucial for understanding its evolution, developing control strategies, and detecting new outbreaks. The easy spread of LSDV through direct contact, vectors, and bodily fluids emphasizes the importance of proper animal management and biosecurity measures to prevent and control the disease.

Sheep poxvirus and Goat poxvirus, both belonging to the genus *Capripoxvirus*, have been endemic in India. However, LSDV outbreaks only started recently in Southeast Asian countries in 2019 [45]. Another poxvirus of the *Poxviridae* family, Smallpox virus, which caused significant damage to human health and life, has been globally eradicated in the last century through large-scale vaccination campaigns using live vaccinia vaccines [46]. Therefore, most effective way to prevent and eliminate disease is vaccination. While there are vaccines for LSD based on the Neethling strain. However, animals sometimes develop clinical symptoms such as the formation of nodules and a drop in milk yield even after vaccination. This adverse effect is known as ‘Neethling disease’ [47]. The commonly used Goat pox vaccine is often employed to prevent LSDV, due to its antigenic similarity to LSDV. The fact that some outbreaks are occurring in vaccinated animals raises about the complete effectiveness of the vaccine. Possible reasons for this include the vaccination method being ineffective, flaws in the administration methods, or issues with vaccine storage.

Some studies report that the commonly used LSDV vaccine is not a pure viral culture of LSDV virus, but instead contains a mix of quasi-species which raises the chances of homologous recombination in the genome of viruses [36]. It has been reported that LSDV can also undergo recombination [31]. Therefore, while using live attenuated viruses, it is important to analyse the quality and nature of variation. Previously, several vaccine-like recombinant LSDV strains were discovered in Kazakhstan, neighbouring of Russia and China from 2017-2019 [48]. In our study, we analysed the current LSDV outbreak clusters and observed that both the groups lie away from the vaccine strains as well as the recombination derived strains, therefore the current outbreak is not a result of vaccine spillover. The 22 LSDV genomes sequenced in this study are separate from the recombinant LSDV strains from Southeast Asia and the Russian vaccine spillover LSDV strains, as clearly depicted in the phylogenetic analysis.

A Ranchi strain based homogenous live-attenuated LSD vaccine, Lumpi-Pro VacInd has been developed after passaging the virus in cell culture towards preventing the outbreaks. The ‘Neethling disease’ event in which animals develop reaction against vaccination such as skin nodule formation was not observed in field animals and the vaccine was shown to be 100% effective till January 2023 [49]. The SNP analysis of this vaccine strain showed that it belongs to the LSDV sequences of Group I in the phylogram which has lesser number of mutations. It is not known if the vaccine will be able to prevent the infection caused by the highly mutating strains in Group II, but it has been shown that it neutralize LSDV strains from 2022 outbreak [49]. Continued monitoring of vaccinated animals for extended periods will be important to assess the vaccine’s efficacy.

To prevent future outbreaks of LSDV, it is crucial to identify the possible reasons behind these outbreaks and strategize accordingly. It is important to identify all potential hosts for LSDV beyond cattle and arthropod vectors. Regulating the transboundary migrations of animals is necessary to prevent all possible pathways for LSDV introduction. Regarding vaccines, live attenuated LSDV strain vaccines should only be used after undergoing quality testing.

## CONCLUSION

The analysis of LSDV cases in India during the 2022 outbreak highlighted the severe impact of the disease on cattle and the significant economic losses incurred in the agricultural sector. Developing a molecular detection strategy using PCR allowed for accurate diagnosis of LSDV, overcoming the challenges posed by clinical signs overlapping with other diseases. Whole genome sequencing provided valuable information on the origin and evolution of the LSDV strains circulating in India. The phylogenetic analysis revealed the presence of two distinct groups, suggesting multiple introductions of the virus. SNP analysis identified genetic variations in essential genes, potentially affecting virulence and antigenicity.

These findings contribute to a better understanding of LSDV and provide valuable information for developing effective control strategies, such as vaccines and diagnostic tests. Monitoring genetic variations and protein expression patterns is crucial for detecting new outbreaks, tracking the virus’s evolution, and guiding the development of targeted interventions. Overall, this research enhances our knowledge of LSDV and its impact on cattle health, supporting efforts to mitigate its spread and minimize economic losses in the future.

## AUTHOR CONTRIBUTIONS

UT^1^, GR^3^, BG^3^ and CJ^7^ conceived and designed the study. PY^1^, AK^1^ performed the ONT based sequencing, analysed the data, and wrote the manuscript. AP^7^, SS^7^, JR^7^, RP^7^, PC^7^ prepared samples and performed NGS Illumina sequencing. MJ^7^, DP^4^, FT^5^, NK^6^ helped in coordinating with team. SN^1^, SN^3^, YR^3^ and RSK^4^ diagnosed the cattle. SN^1^, JR, AK, NK collected samples from field. UT^1^, GR^3^, BG^3^, CJ^7^ reviewed the manuscript.

## DATA AVAILABILITY STATEMENT

The data that supports the findings of this study is available in the NCBI (BioProject ID PRJNA998208 and PRJNA1004082). The fasta sequences for the isolates reported in the study are available at doi.org/10.6084/m9.figshare.24018948.

## REFERENCES

1. Alexander R, Plowright W, Haig D (1957) Cytopathogenic agents associated with lumpy skin disease of cattle. Bulletin of epizootic diseases of Africa 5: 489–492.

2. Diallo A, Viljoen GJ (2007) Genus Capripoxvirus. In: Mercer AA, Schmidt A, Weber O, editors. Poxviruses. Basel: Birkhäuser Basel. pp. 167–181.

3. Sanz-Bernardo B, Haga IR, Wijesiriwardana N, Hawes PC, Simpson J, et al. (2020) Lumpy Skin Disease Is Characterized by Severe Multifocal Dermatitis With Necrotizing Fibrinoid Vasculitis Following Experimental Infection. Vet Pathol 57: 388–396.

4. Davies FG (1991) Lumpy skin disease, an African capripox virus disease of cattle. Br Vet J 147: 489–503.

5. Babiuk S, Bowden TR, Boyle DB, Wallace DB, Kitching RP (2008) Capripoxviruses: an emerging worldwide threat to sheep, goats and cattle. Transbound Emerg Dis 55: 263–272.

6. Davies FG (1991) Lumpy skin disease of cattle: a growing problem in Africa and the Near East. World Animal Review 68: 37–42.

7. Young E, Basson PA, Weiss KE (1970) Experimental infection of game animals with lumpy skin disease virus (prototype strain Neethling). Onderstepoort J Vet Res 37: 79–87.

8. Dao TD, Tran LH, Nguyen HD, Hoang TT, Nguyen GH, et al. (2022) Characterization of Lumpy skin disease virus isolated from a giraffe in Vietnam. Transbound Emerg Dis 69: e3268–e3272.

9. Kumar R, Godara B, Chander Y, Kachhawa JP, Dedar RK, et al. (2023) Evidence of lumpy skin disease virus infection in camels. Acta Trop 242: 7.

10. Chihota CM, Rennie LF, Kitching RP, Mellor PS (2001) Mechanical transmission of lumpy skin disease virus by Aedes aegypti (Diptera: Culicidae). Epidemiol Infect 126: 317–321.

11. Carn VM, Kitching RP (1995) An investigation of possible routes of transmission of lumpy skin disease virus (Neethling). Epidemiol Infect 114: 219–226.

12. Chihota CM, Rennie LF, Kitching RP, Mellor PS (2003) Attempted mechanical transmission of lumpy skin disease virus by biting insects. Med Vet Entomol 17: 294–300.

13. Tuppurainen ES, Venter EH, Coetzer JA, Bell-Sakyi L (2015) Lumpy skin disease: attempted propagation in tick cell lines and presence of viral DNA in field ticks collected from naturally-infected cattle. Ticks Tick Borne Dis 6: 134–140.

14. Gubbins S (2019) Using the basic reproduction number to assess the risk of transmission of lumpy skin disease virus by biting insects. Transbound Emerg Dis 66: 1873–1883.

15. Weiss K (1968) Lumpy skin disease virus. Cytomegaloviruses Rinderpest Virus Lumpy Skin Disease Virus: Springer. pp. 111–131.

16. Kumar N, Tripathi BN A serious skin virus epidemic sweeping through the Indian subcontinent is a threat to the livelihood of farmers: Virulence. 2022 Dec;13(1):1943–1944. doi: 10.1080/21505594.2022.2141971.

17. Department SR (Dec 1, 2022) Cattle population in India 2016-2023.

18. Ali AA, Neamat-Allah ANF, Sheire HAE, Mohamed RI (2021) Prevalence, intensity, and impacts of non-cutaneous lesions of lumpy skin disease among some infected cattle flocks in Nile Delta governorates, Egypt. Comp Clin Path 30: 693–700.

19. Diseases OO-L (2010) Version adopted by the World Assembly of Delegates of the OIE in May 2010, OIE, Paris.: Terrestrial Manual of Lumpy Skin Disease.

20. MacOwan K (1959) Observations on the epizootiology of lumpy skin disease during the first year of its occurrence in Kenya. Bull Epizootic Dis of Africa 7: 7–20.

21. Tuppurainen ES, Oura CA (2012) Review: lumpy skin disease: an emerging threat to Europe, the Middle East and Asia. Transbound Emerg Dis 59: 40–48.

22. House JA, Wilson TM, Nakashly SE, Karim IA, Ismail I, et al. (1990) The isolation of lumpy skin disease virus and bovine herpesvirus-from cattle in Egypt. Journal of Veterinary Diagnostic Investigation 2: 111–115.

23. Lojkić I, Šimić I, Krešić N, Bedeković T (2018) Complete genome sequence of a lumpy skin disease virus strain isolated from the skin of a vaccinated animal. Genome Announc 6: e00482–00418.

24. Kumar N, Chander Y, Kumar R, Khandelwal N, Riyesh T, et al. (2021) Isolation and characterization of lumpy skin disease virus from cattle in India. PLoS One 16: e0241022.

25. Salib FA, Osman AH (2011) Incidence of lumpy skin disease among Egyptian cattle in Giza Governorate, Egypt. Veterinary world 4.

26. Babiuk S, Bowden T, Parkyn G, Dalman B, Manning L, et al. (2008) Quantification of lumpy skin disease virus following experimental infection in cattle. Transboundary and Emerging diseases 55: 299–307.

27. Davies F (1991) Lumpy skin disease, an African capripox virus disease of cattle. British Veterinary Journal 147: 489–503.

28. Irons P, Tuppurainen E, Venter E (2005) Excretion of lumpy skin disease virus in bull semen. Theriogenology 63: 1290–1297.

29. Mathivanan EME, Raju K, Murugan R (2023) Outbreak of Lumpy skin disease in India 2022 - an emerging threat to livestock & livelihoods. Global Biosecurity 5.

30. Sudhakar SB, Mishra N, Kalaiyarasu S, Jhade SK, Hemadri D, et al. (2020) Lumpy skin disease (LSD) outbreaks in cattle in Odisha state, India in August 2019: Epidemiological features and molecular studies. Transboundary and Emerging diseases 67: 2408–2422.

31. Minor PD, John A, Ferguson M, Icenogle JP (1986) Antigenic and molecular evolution of the vaccine strain of type 3 poliovirus during the period of excretion by a primary vaccinee. Journal of general virology 67: 693–706.

32. K LK, B S, K R (2021) Clinico-molecular diagnosis and characterization of bovine lumpy skin disease virus in Andhra Pradesh, India. Tropical Animal Health and Production 53: 424.

33. Quick J, Grubaugh ND, Pullan ST, Claro IM, Smith AD, et al. (2017) Multiplex PCR method for MinION and Illumina sequencing of Zika and other virus genomes directly from clinical samples. Nat Protoc 12: 1261–1276.

34. Whitford W, Hawkins V, Moodley KS, Grant MJ, Lehnert K, et al. (2022) Proof of concept for multiplex amplicon sequencing for mutation identification using the MinION nanopore sequencer. Sci Rep 12: 022–12613.

35. Besemer J, Lomsadze A, Borodovsky M (2001) GeneMarkS: a self-training method for prediction of gene starts in microbial genomes. Implications for finding sequence motifs in regulatory regions. Nucleic Acids Res 29: 2607–2618.

36. Sprygin A, Pestova Y, Bjadovskaya O, Prutnikov P, Zinyakov N, et al. (2020) Evidence of recombination of vaccine strains of lumpy skin disease virus with field strains, causing disease. PLoS One 15.

37. Chatterjee M, Osborne J, Bestetti G, Chang Y, Moore PS (2002) Viral IL-6-induced cell proliferation and immune evasion of interferon activity. Science 298: 1432–1435.

38. Tulman ER, Afonso CL, Lu Z, Zsak L, Kutish GF, et al. (2001) Genome of lumpy skin disease virus. J Virol 75: 7122–7130.

39. Van Vliet K, Mohamed MR, Zhang L, Villa NY, Werden SJ, et al. (2009) Poxvirus proteomics and virus-host protein interactions. Microbiol Mol Biol Rev 73: 730–749.

40. Moon KB, Turner PC, Moyer RW (1999) SPI-1-dependent host range of rabbitpox virus and complex formation with cathepsin G is associated with serpin motifs. J Virol 73: 8999–9010.

41. Chibssa TR, Sombo M, Lichoti JK, Adam TIB, Liu Y, et al. (2021) Molecular Analysis of East African Lumpy Skin Disease Viruses Reveals a Mixed Isolate with Features of Both Vaccine and Field Isolates. Microorganisms 9: 1142.

42. Babkin I, Shchelkunov S (2006) Time scale of poxvirus evolution. Molecular Biology 40: 16–19.

43. Van Schalkwyk A, Byadovskaya O, Shumilova I, Wallace DB, Sprygin A (2022) Estimating evolutionary changes between highly passaged and original parental lumpy skin disease virus strains. Transbound Emerg Dis 69: e486–e496.

44. Sudhakar SB, Mishra N, Kalaiyarasu S, Ahirwar K, Chatterji S, et al. (2023) Lumpy Skin Disease Virus Infection in Free-Ranging Indian Gazelles (Gazella bennettii), Rajasthan, India. Emerg Infect Dis 29: 1407–1410.

45. Das M, Chowdhury MSR, Akter S, Mondal AK, Uddin M, et al. (2021) An updated review on lumpy skin disease: Perspective of southeast asian countries. J Adv Biotechnol Exp Ther 4: 322–333.

46. Strassburg MA (1982) The global eradication of smallpox. Am J Infect Control 10: 53–59.

47. Ben-Gera J, Klement E, Khinich E, Stram Y, Shpigel NY (2015) Comparison of the efficacy of Neethling lumpy skin disease virus and x10RM65 sheep-pox live attenuated vaccines for the prevention of lumpy skin disease - The results of a randomized controlled field study. Vaccine 33: 4837–4842.

48. Vandenbussche F, Mathijs E, Philips W, Saduakassova M, De Leeuw I, et al. (2022) Recombinant LSDV Strains in Asia: Vaccine Spillover or Natural Emergence? Viruses 14.

49. Kumar N, Barua S, Kumar R, Khandelwal N, Kumar A, et al. (2023) Evaluation of the safety, immunogenicity and efficacy of a new live-attenuated lumpy skin disease vaccine in India. Virulence 14: 2190647.

